# MetaLab Platform Enables Comprehensive DDA and DIA Metaproteomics Analysis

**DOI:** 10.1101/2024.09.27.615406

**Authors:** Kai Cheng, Zhibin Ning, Xu Zhang, Haonan Duan, Janice Mayne, Daniel Figeys

**Affiliations:** School of Pharmaceutical Sciences, Faculty of Medicine, University of Ottawa, Ottawa, Canada; Regulatory Research Division, Biologic and Radiopharmaceutical Drugs Directorate, Health Products and Food Branch, Health Canada, Ottawa, Canada

**Keywords:** metaproteomics, gut microbiome, mass spectrometry, DIA, software

## Abstract

Metaproteomics studies the collective protein composition of complex microbial communities, providing insights into microbial roles in various environments. Despite its importance, metaproteomic data analysis is challenging due to the data’s large and heterogeneous nature. While Data-Independent Acquisition (DIA) mode enhances proteomics sensitivity, it traditionally requires Data-Dependent Acquisition (DDA) results to build the library for peptide identification.

This paper introduces an updated version of MetaLab, a software solution that streamlines metaproteomic analysis by supporting both DDA and DIA modes across various mass spectrometry (MS) platforms, including Orbitrap and timsTOF. MetaLab’s key feature is its ability to perform DIA analysis without DDA results, allowing more experimental flexibility. It incorporates a deep learning strategy to train a neural network model, enhancing the accuracy and coverage of DIA results.

Evaluations using diverse datasets demonstrate MetaLab’s robust performance in accuracy and sensitivity. Benchmarks from large-scale human gut microbiome studies show that MetaLab increases peptide identification by 2.7 times compared to conventional methods. MetaLab is a versatile tool that facilitates comprehensive and flexible metaproteomic data analysis, aiding researchers in exploring microbial communities’ functionality and dynamics.

## Main

Microbial communities are intricate networks of microorganisms that inhabit various environments, playing critical roles in ecosystem functions such as nutrient cycling, disease resistance, and overall ecosystem stability. Understanding the functional dynamics of these communities is essential for revealing their roles in ecosystem processes and for advancing applications in biotechnology, agriculture, and human health^1–3^. Metaproteomics has emerged as a powerful approach to studying these microbial communities by examining their protein profiles^4^. By characterizing the proteins expressed within these communities, metaproteomics offers valuable insights into their metabolic activities, interactions, and adaptive responses to environmental changes. However, the analysis of metaproteomic data presents significant challenges due to the inherent complexity and heterogeneity of microbial communities and the vast diversity of their expressed proteins^5^.

Two widely used approaches in mass spectrometry-based proteomics are Data-Dependent Acquisition (DDA) and Data-Independent Acquisition (DIA)^6^. DDA typically involves selecting and fragmenting the most abundant precursor ions for analysis, offering high accuracy but potentially underrepresenting low-abundance peptides. In contrast, DIA involves fragmenting all ions within a defined mass range, providing a comprehensive view of the proteome but presenting challenges for data interpretation. The limitations of current metaproteomic data analysis strategies become increasingly apparent when applied to DIA data. Most DIA data analysis approaches rely on spectrum/peptide libraries built from DDA datasets, which is time-consuming and labor-intensive^7^. Moreover, DIA identifications are restricted to DDA results, limiting the advantages of DIA sensitivity and comprehensive proteome coverage. *GlaDIator*^8^ introduced a DIA-only method that leverages the DIA-Umpire^9^ algorithm to deconvolve DIA spectra into pseudo-DDA spectra for subsequent database searches. However, recent advancements in efficient software and algorithms, such as Dia-NN^10^ and Spectronaut®, have made direct analysis of DIA spectra a more favored approach. This approach offers improved accuracy, sensitivity, and computational efficiency compared to the traditional pseudo-MS/MS spectrum generation method.

Our previous efforts introduced MetaLab^11^ and MetaLab-MAG^12^ for the deep characterization of DDA data from various microbiota samples, utilizing strategies such as searching against integrated gene catalogue databases^13^ and biome-specific microbial genome catalogues^14^, respectively. The MetaLab-MAG workflow proved advantageous, enabling targeted analysis limited to specific microbiomes, resulting in more relevant and accurate results compared to broader gene catalog databases, and improving the resolution of taxonomic analysis.

This paper introduces updated versions of MetaLab-MAG and MetaLab-DIA. MetaLab-MAG now includes capabilities for analyzing timsTOF datasets, leveraging the benefits of timsTOF technology, such as improved ion mobility separation and higher sensitivity for metaproteomic studies. Furthermore, we introduce a novel scoring scheme for evaluating genome-level identifications, enhancing the accuracy and reliability of metaproteomic data analysis.

MetaLab-DIA, tailored for metaproteomics DIA data analysis, utilizes a neural network model built from prior quantitative results to predict peptide candidates for DIA identification. The workflow, like MetaLab-MAG, involves determining possible genome candidates first and selecting peptides with higher predicted intensities to build the library for DIA search. This method doubled peptide identifications in DIA mode compared to DDA mode across multiple datasets, a result unachievable by conventional DDA-based DIA searches. MetaLab-DIA overcomes the limitations of existing approaches, providing researchers with a comprehensive solution for metaproteomic data analysis. Specifically, we present 15 human gut microbiota models derived from healthy, inflammatory bowel disease, and Type 1 diabetes samples, offering researchers greater flexibility and efficiency in exploring metaproteomic datasets (**Supplementary note**).

In summary, the MetaLab software suite seamlessly supports both DDA and DIA modes on diverse mass spectrometry (MS) platforms, including high-resolution Orbitrap and timsTOF instruments. By integrating these platforms and introducing innovative data analysis approaches, MetaLab enhances metaproteomic analyses’ depth and accuracy, offering a comprehensive solution to the challenges posed by microbial community proteome studies.

## Results

### Overview of MetaLab Software

The new version of MetaLab is a comprehensive software suite designed for metaproteomics data analysis, supporting both DDA and DIA modes across multiple MS platforms. The previous version of MetaLab-MAG utilized the pFind search engine due to its high identification rate^15^. However, it lacked native support for .d spectra format. To overcome this limitation, we integrated FragPipe and AlphaPept, both of which provide native support for .d datasets^16–18^.

MetaLab-MAG’s iterative search strategy begins by determining a list of genome candidates, followed by extracting all proteins from these genomes to create a combined database for subsequent searches. This strategy is also applicable to DIA data analysis by incorporating the DIA data search engine DIA-NN^10^ into the workflow (**Figure 1a**). However, the performance of DIA database searches is more sensitive to database size compared to DDA. Simply combining all proteins from candidate genomes into a sample-specific database may not be optimal for DIA searches.

**Figure 1.**
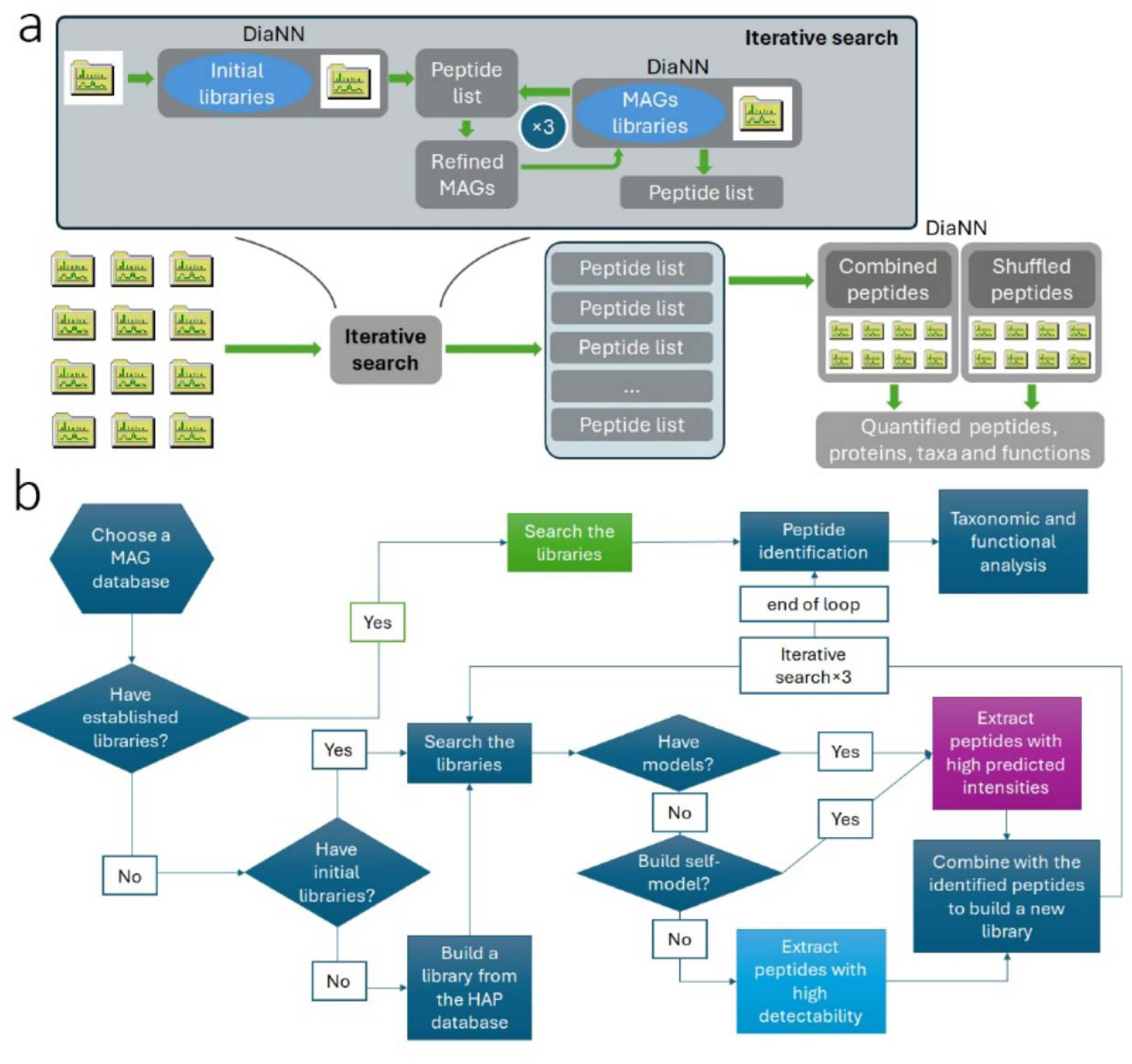
(a) overview of computational workflow of MetaLab-DIA. (b) the diverse strategies of the iterative search workflows.

To address this challenge, we propose diverse solutions based on available resources (**Figure 1b**). If a complete and properly sized library is available, users can search this library to obtain taxonomic and functional analysis based on peptide identification. If this resource is absent, the iterative search begins using an initial library derived from a peptide/protein sequence list related to the sample, such as database search results from similar samples. The raw files are searched against this initial library to generate a peptide list that determines the genome candidates. If peptide/protein identification results are not available, a high-abundance protein database can be used as an alternative.

After determining the genome candidates, instead of using all in silico digested peptides from these genomes, only peptides more likely to be identified are selected to form the library for the iterative search. This selection is the key step of MetaLab-DIA, with two provided options. The first is a general solution using peptides with high predicted detectability^19^. The second utilizes peptide quantitative information to build a deep learning model. Quantitative information can come from three sources: 1) publicly available datasets from similar samples; 2) prior experimental results from the same sample; 3) this dataset itself. This model predicts the intensity of the in silico digested peptide, enabling the selection of peptides with high predicted intensity to form the library for DIA search. This strategy incorporates comprehensive peptide information beyond detectability, including biome-specific factors such as genome, taxon, and function abundance. The second strategy is recommended and explained in detail below.

The user specifies a genome catalogue, with MetaLab currently supporting 11 types of genome catalogues from MGnify^20, 21^. All proteins from the genome catalogue are in silico digested into peptides, and the peptide detectability is predicted based on peptide sequence properties. To build a prediction model, a peptide list with intensity information is required, such as the DDA quantitative result of the same sample or publicly available datasets from similar samples. If none are available, a pre-search of this dataset itself can provide the quantitative information. This peptide list is mapped to in silico digested peptides and linked to corresponding proteins from the genome catalogue. Based on protein information, intensities of corresponding genomes, taxa, and functions are obtained.

Consequently, a data table is created, encompassing 16 features including peptide intensity, peptide detectability, protein intensity, genome intensity, taxa intensities at seven levels, and function intensities at five levels. This table serves as the training dataset for building a model to predict peptide intensity, employing Deep Feedforward Neural Network (DFNN) or Recurrent Neural Network (RNN) learning methods. RNN performs better on our test data, so it is set as the default method. Once trained, the model predicts peptide intensities within the specified genome catalogue. These peptides are then ranked based on predicted intensities, forming the primary source for the iterative search process.

After the first search, a candidate genome list is obtained for each raw file, from which high-ranked peptides are selected to compose a peptide library, limited to two million peptides. This process is repeated, and after three search loops, a peptide list result is obtained for each raw file. These lists are combined to form a complete peptide library for the final search of all raw files in the task. Identified peptides and their intensities from the final search are used for taxonomic and functional analysis.

### Improved Metaproteomic Analysis with MetaLab-MAG and MetaLab-DIA

We utilized two distinct datasets for the evaluation: microbial mixtures containing twelve different bacterial strains (referred to as 12mix) and human fecal samples from Aakko’s paper^22^. These samples were analyzed using both DDA and DIA modes. For the 12mix, each mode was run with three technical replicates, while for the complex human fecal samples, six technical replicates were performed per mode.

For the DDA datasets, the raw files were directly searched against the Integrated Gene Catalog (IGC) database used in the paper. Due to the vast database size, database searches resulted in low identification rates, yielding 20,047 and 24,839 peptide sequences from the two datasets, respectively. Using the MetaLab-MAG pFind workflow, peptide identification rates were significantly improved, resulting in 36,900 and 38,017 peptide sequences, representing increases of 84.1% and 53.1%, respectively (**Figure 2a-b, Supplementary Data 1, 2**).

**Figure 2.**
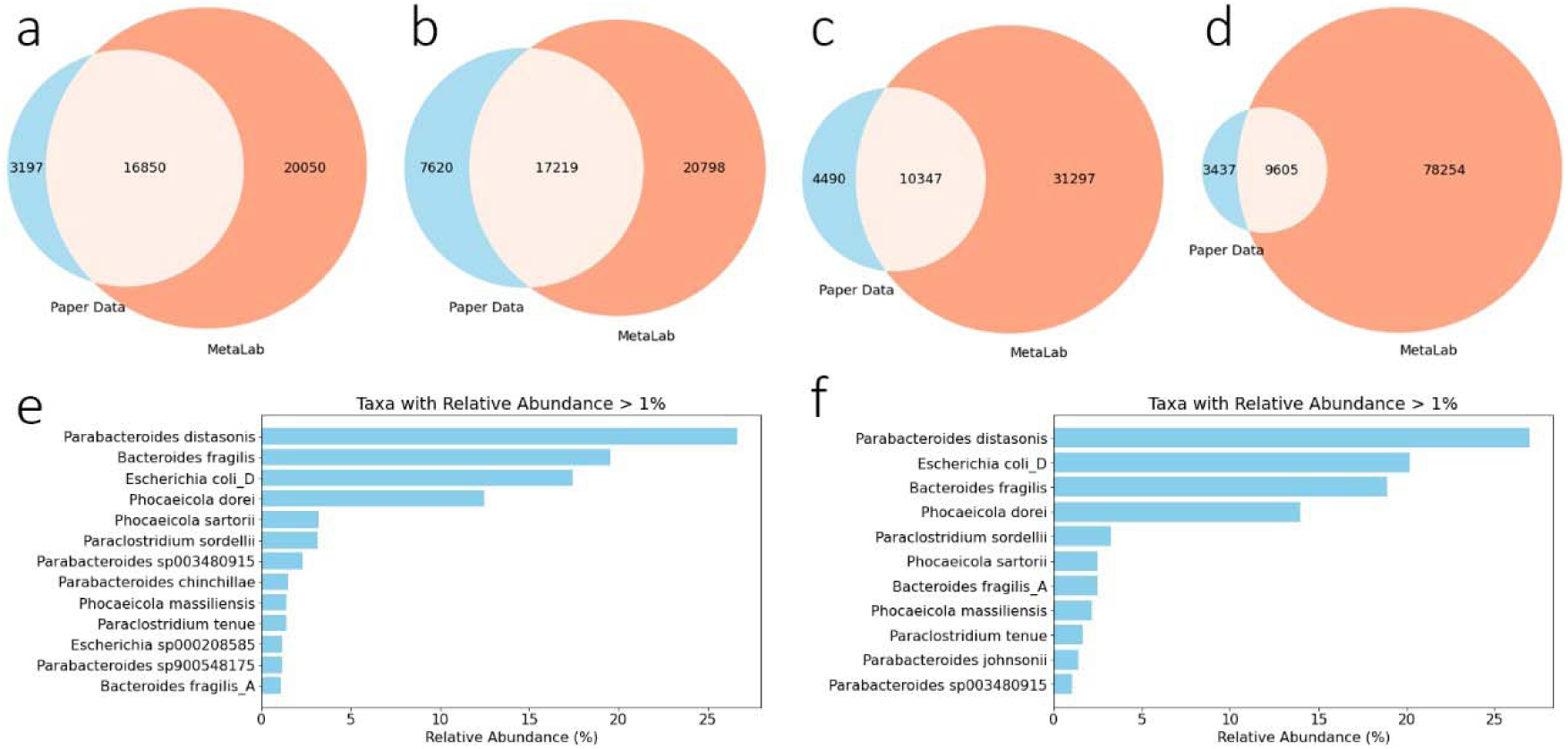
**(a)-(d)** the overlap between the peptides identified by the original paper and our strategy. **(a)** 12mix DDA dataset; **(b)** fecal DDA dataset; **(c)** 12mix DIA dataset; **(d**) fecal DIA dataset. **(e)** the high-abundance species in the 12mix DDA dataset and **(f)** 12mix DIA dataset.

In Aakko’s paper, the DIA data was analyzed using the conventional DDA-based strategy, that is, searched against the spectral library derived from the DDA datasets. The number of peptide identifications was 14,837 and 13,042 from the 12mix and fecal samples, accounting for 74.0% and 52.5% of the DDA peptide identifications, respectively. This demonstrated that the more complex the sample, the lower the proportion of peptides in the DDA-based library that could be identified from DIA datasets. Consequently, the DDA analysis could not characterize the sample with sufficient depth, making the DDA-based library deficient for comprehensive DIA mode analysis. To address this, they developed *GlaDIator* which can analyze the DIA data directly by converting the DIA spectra to pseudo-DDA spectra. Through this strategy, 7,967 and 14,691 peptides were identified^8^.

In contrast, by utilizing a DDA-based initial library and model, the MetaLab-DIA workflow identified 41,881 and 94,951 peptides from the two samples, respectively (**Figure 2c-d, Supplementary Data 3, 4**). Compared to the paper’s results, this represents increases of 2.8 and 7.3 times. Compared to the DDA results obtained using MetaLab-MAG, DIA mode resulted in 114% and 250% more peptide identifications. Notably, more complex fecal samples showed a greater increase in identified peptides in DIA than in DDA mode, clearly demonstrating that the MetaLab-DIA strategy leveraged the high sensitivity of DIA mode.

The credibility of peptide identifications from the 12mix samples was evaluated by examining the diverse array of bacteria, which included 10 different genera. Among these genera, eight were identified by MetaLab-MAG, except for *Streptococcus* and *Enterorhabdus* (the latter was not included in the MGnify database). Remarkably, 97.7% and 98.1% of the peptides derived from the 12mix DDA and DIA datasets were attributed to the correct eight genera, underscoring the reliability of the peptide identifications achieved with MetaLab. Furthermore, MetaLab introduced a scoring scheme to evaluate identified genomes. Applying this scheme to the 12mix dataset, we identified 45 and 33 genomes from DDA and DIA data, respectively, with a p-value less than 0.01, of which 25 and 26 were associated with the correct genera. At the species level, all species with relative abundance above 1% from the two datasets were from the correct genera (**Figure 2e, 2f**), highlighting the confidence of the identification in genome level.

### Analyzing timsTOF Data from Mouse Gut Microbiome Samples

We evaluated MetaLab’s performance using a mouse gut microbiome dataset from the timsTOF platform^23^. This dataset was generated using both DDA- and DIA-PASEF methods, with 30 raw files collected for each mode.

For the DDA-PASEF dataset, we employed the FragPipe workflow in MetaLab-MAG version. The database used was the Mouse Gut v1.0 genome catalogue from MGnify, containing 2,847 species. The range of PSMs (Peptide-Spectrum Matches) and peptides identified from a single raw file in the DDA dataset were 33,849±6,800 (mean ± standard deviation, same below) and 14,992±3,216, respectively (**Figure 3a-b, Supplementary Data 5**). The total number of peptide and protein identifications were 53,848 and 12,250, respectively. We then built a prediction model based on this peptide list and performed the DIA-PASEF data analysis using MetaLab-DIA. On average, 39,833±8,666 PSMs and 37,603±7,781 peptides per raw file were obtained (**Figure 3a-b, Supplementary Data 6**). The total number of peptide and protein identifications were 66,259 and 17,626, respectively.

**Figure 3.**
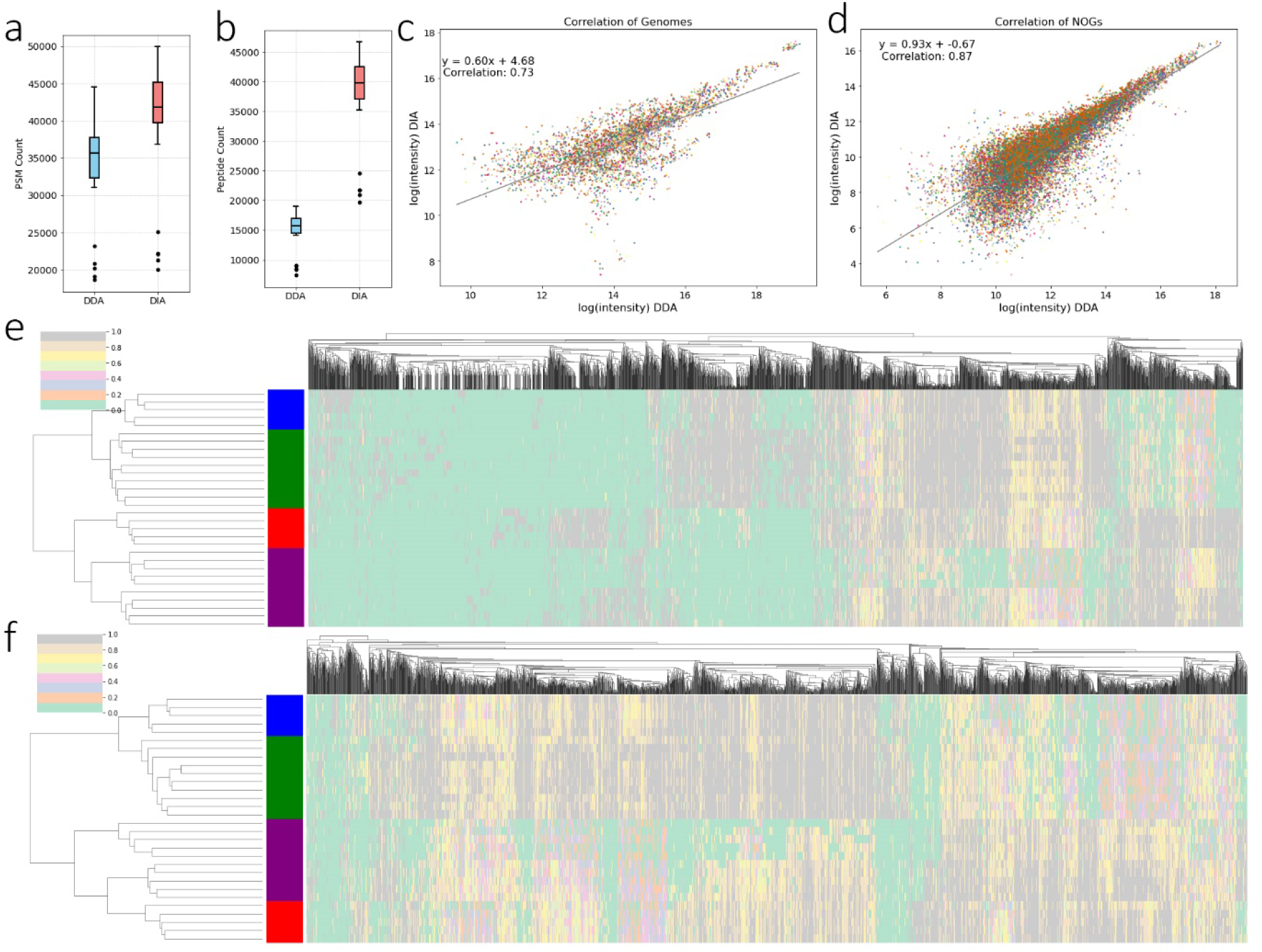
the comparison between the results from DDA and DIA datasets. **(a**) PSM count; **(b)** peptide count; **(c)** the correlation of quantified genomes; **(d)** the correlation of quantified NOGs; **(e)** the cluster of the NOGs from DDA result; **(f)** the cluster of the NOGs from DIA result. Blue: differential centrifugation then in-solution digestion; green: differential centrifugation then Filter-Aided Sample Preparation (FASP) digestion; red: non-differential centrifugation then in-solution digestion; purple: non-differential centrifugation then FASP digestion.

The DIA dataset generated 2.5 times more peptides per raw file and 23.0% more peptide identifications for the entire dataset compared to the DDA. Notably, these statistics reflect modified peptides; considering only the peptide sequences, 33.5% more peptide sequences were identified from the DIA dataset. This data demonstrates that DIA has greater depth than DDA. Consequently, it is limiting to directly use the spectral library obtained from DDA data for DIA data retrieval. However, using the model derived from the DDA results, we can enhance the spectral library for DIA data analysis by including peptide candidates not directly identified by DDA but likely present in the sample.

From the DIA dataset, we identified 185 genomes and 17,505 functions. The quantitative results at the genome level showed high consistency between the DDA and DIA results (**Figure 3c**). In contrast, at the function level, the consistency was even higher, with a correlation coefficient of 0.87 between DDA and DIA results (**Figure 3d**). This indicates that despite slight differences in species quantified by different methods, the functional quantifications remain highly consistent. This high consistency also reflects the reliability of quantitative results from both methods. This conclusion is further supported by the clustering results (**Figures 3e-f**), where samples generated from different sample preparation methods clustered differently based on quantitative information at the function level. Although the results appear similar, we observed a significant advantage of the DIA approach with greatly reduced occurrence of missing values in the samples, which significantly aids in studying quantitative changes in specific functions.

### Analyzing Orbitrap Data from Human Gut Microbiome Samples

To evaluate DDA and DIA performance in human gut microbiome analysis, we tested one sample with a high degree of technical duplication^24^. In both modes, 21 technical replicates were conducted on the QE platform. For the DDA dataset, an open search strategy enabled comprehensive post-translational modification identifications. In this case, 55,877 peptides from 42,539 peptide sequences were identified (**Supplementary Data 7**). From the DIA dataset, 43,542 peptide sequences were identified using the MetaLab-DIA iterative search workflow, which utilized the DDA quantitative result as the training dataset to build the model (**Supplementary Data 8**). Both methods have their advantages: DIA identified more peptide sequences, while DDA identified more modified peptide forms.

Additionally, we used a DDA-independent workflow that employed a core peptide library^25^ (termed MetaPep library) and six trained models based on publicly available datasets^26–30^. In this case, the raw files were searched against the MetaPep library first, to determine the candidate genomes. Then peptides from the candidate genomes with higher predicted intensities by these six models were selected to form the library for the iterative search. This approach identified 48,368 peptides, the greatest number of peptide identifications among the three workflows (**Supplementary Data 9**). Compared to the total number of identifications, the increase in the number of peptides identified from each file was more pronounced in DIA mode. As shown in **Figure 4a**, a greater number of PSMs were identified by the DDA mode. However, the number of peptides identified in DIA mode was approximately 1.5 times that of DDA mode (**Figure 4b**), indicating that DIA mode enabled deeper characterization of the samples than DDA.

**Figure 4.**
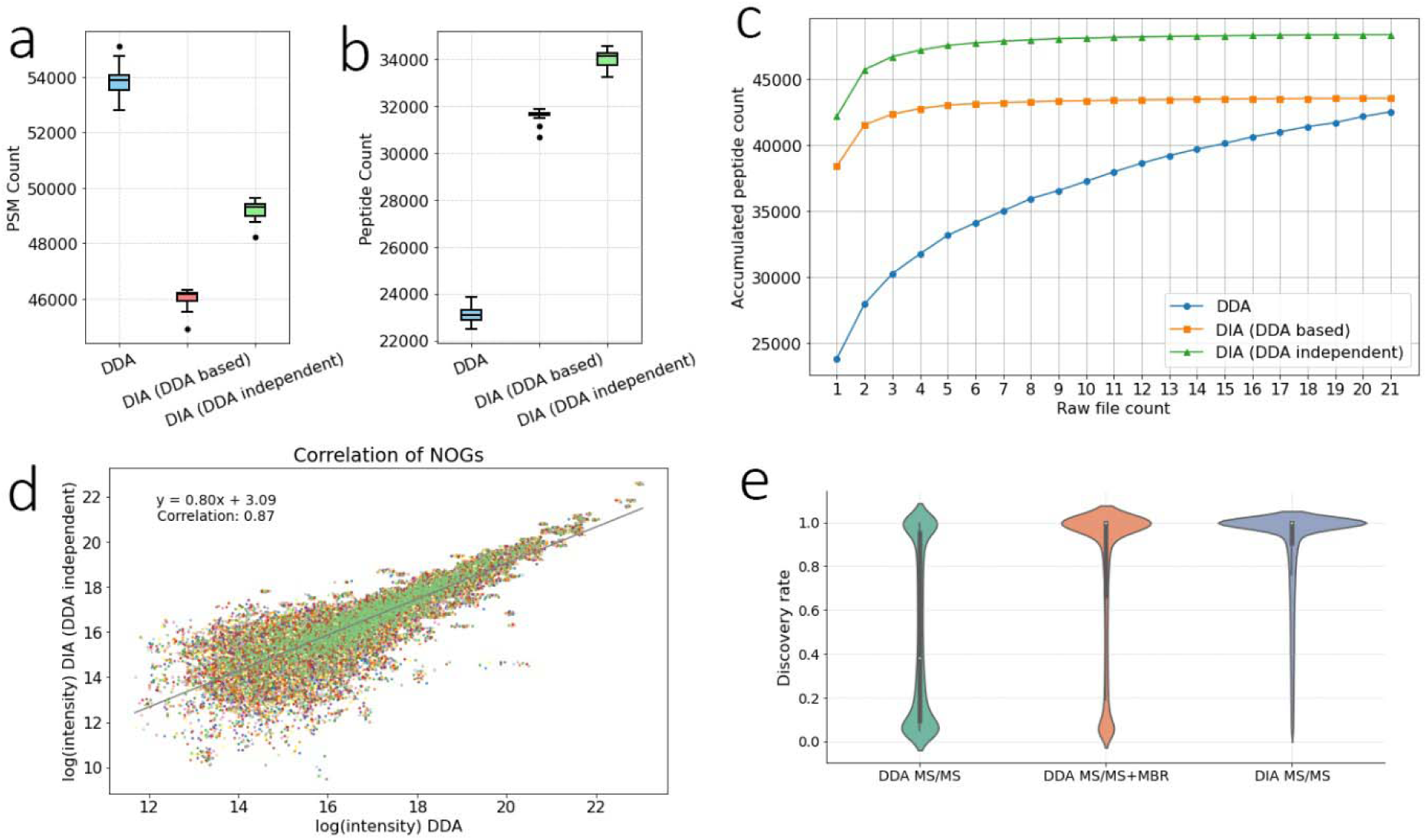
the comparison of the performance crosses the three methods. DDA: DDA datasets; DIA (DDA-based): searching the DDA-derived library and using DDA-based model to predict the peptide candidates; DIA (DDA-independent): searching the library and using the model built by the public available datasets. **(a)** the comparison of the PSM counts. **(b)** the comparison of the peptide counts. **(c)** the comparison of the accumulated peptide counts. **(d)** the comparison of the discovery rate. **(e)** the correlation of NOGs between DDA and DIA by DDA-independent method.

Figure 4c demonstrates that even after 21 technical replicates, the number of accumulated peptides continued to grow in DDA mode, despite the 21 technical replicates are from the same microbiome. It is known that microbiome compositions differ between individuals, which would make getting a comprehensive peptide lists by DDA very demanding. In contrast, the number of accumulated peptides in DIA mode reached an upper limit quickly. We also compared the quantitative results between DDA and DIA modes at the function levels. The correlation between these two modes was 0.87 (Figure 4d). In agreement with the other datasets, this analysis again demonstrated that DDA and DIA showed higher consistency at the functional level, proving that both quantitative methods are reliable.

In addition to accuracy in quantitative analysis, the number of missing values also affects the quantitative outcome. In this dataset, we examined the missing values for both methods. The distribution of discovery rates is illustrated in Figure 4e. For the DDA dataset, if the match-between-run (MBR) strategy was not utilized, the discovery rate of most peptides was below 0.5. This situation improved significantly after using MBR, but the performance was still not as good as in DIA mode. In DIA mode, most peptides had a discovery rate of 1.0, indicating they were detected in all 21 samples. It is worth noting that the MBR strategy only utilized information from MS1, specifically the mass-to-charge ratio of the precursor ion and the retention time. Coeluting ions can affect the accuracy of quantitation, which is more common with complex samples. In DIA mode, not only is the information from MS1 used, but also the fragment ion information from MS2. This allows the DIA mode to achieve a higher discovery rate and more accurate quantification.

In this dataset, the DIA-independent workflow performed even better than the DDA-based workflow, suggesting that the combination of multiple models from public datasets is sufficient for the characterization of common human gut microbiome samples. Consequently, a sample-specific library built from DDA results is not required.

### Large-Scale Human Gut Microbiome Dataset Analysis

To further demonstrate the robustness and scalability of MetaLab-DIA, we analyzed a large human gut microbiome dataset from the timsTOF platform, consisting of 81 raw files^31^. This comprehensive collection of human gut microbiome data provided an unprecedented opportunity to validate MetaLab-DIA’s capabilities on a large scale.

For this analysis, we utilized a DDA-independent method. Instead of using existing models from publicly available DDA results, we built models based on this dataset itself. Initially, all raw files were searched against the MetaPep library. Quantitative information of peptides from each raw file was collected, and the correlation between files was calculated. Rather than using a uniform model, the raw files were grouped based on correlation, resulting in 76 groups. For each group, an independent model was built.

Each iterative search step used peptides with higher intensity predicted by the corresponding model to form sample-specific databases for peptide identification.

From the MetaPep library, we identified an average of 48,640 ± 14,445 peptides per raw file (Figure 5a). Comparatively, searching the DDA-derived database yielded 56,576 ± 17,410 peptides, showing better performance than the MetaPep library in this initial step. However, in the subsequent round, searching the predicted peptide library identified 61,898 ± 16,762 peptides. The cumulative peptide counts from these two steps were 63,405 ± 18,731 (Figure 5a). The iterative search continued twice more to optimize peptide candidates for each raw file, resulting in an accumulated peptide count of 68,260 ± 20,161 (Figure 5a). The combined peptide library size reached 760,201, approximately 2.7 times larger than the DDA-derived database (285,283 peptides). Ultimately, the DDA-independent strategy identified 88,889 ± 28,328 peptides per raw file (Figure 5a**, Supplementary Data 10**), with a total of 559,572 peptides quantitatively identified. In comparison, only 228,746 peptides were identified using the DDA-derived database (Figure 5b), demonstrating a 2.4-fold improvement with our strategy.

**Figure 5.**
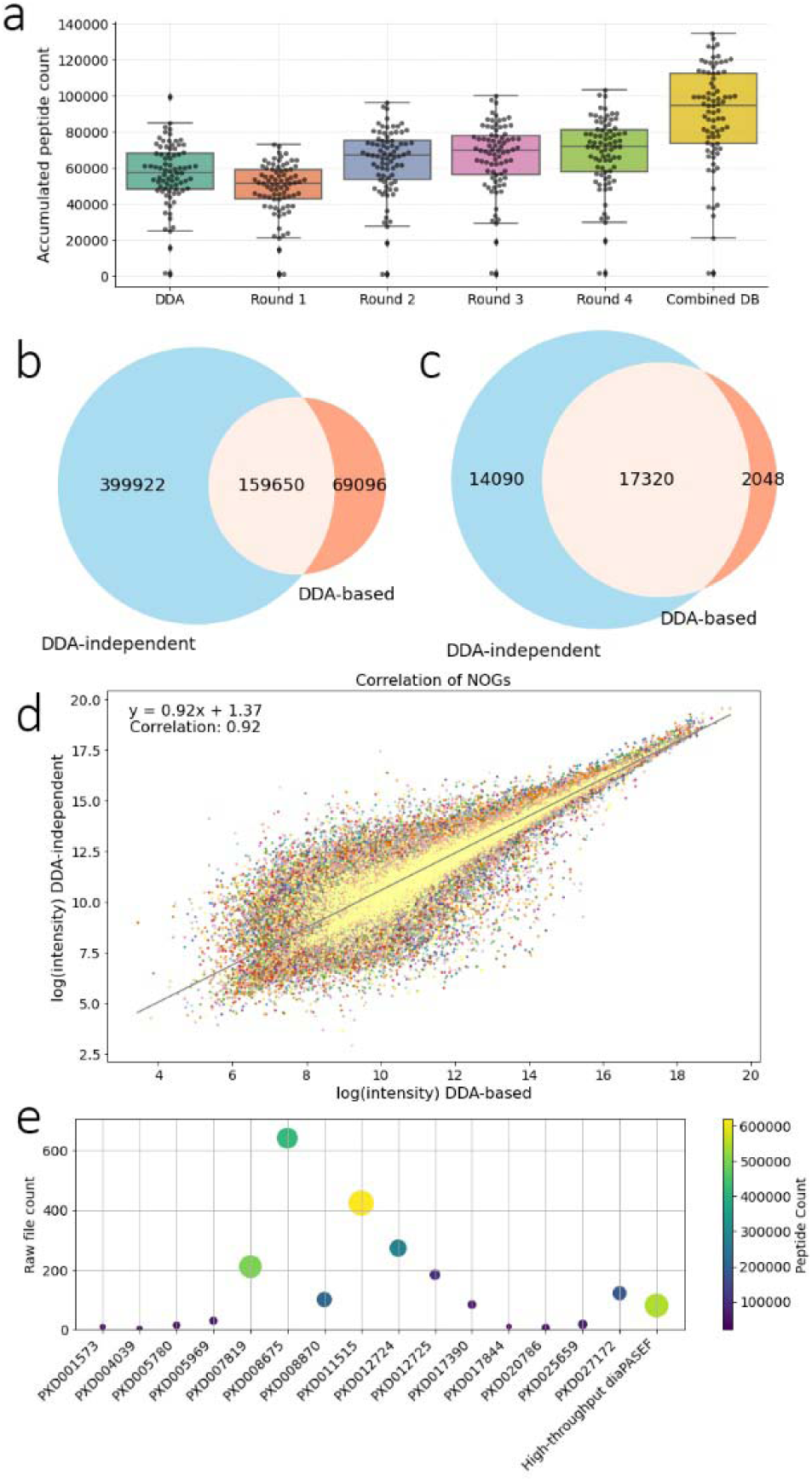
**(a)** the comparison of the peptide count distributions cross different methods/steps. DDA: searching against the DDA-derived peptide library; Round 1: searching against the MetaPep library; Round 2-4: searching against the peptide library based on the model predicted peptides with higher intensities. Please notice the Y-axis shows the accumulated peptide count, which means Round 2 represents unique peptide sequence from both Round 1 and Round 2, and so on. Combined DB: the peptide identified from the combined database composing by the peptide list from each raw file. **(b)-(c)** the overlap of (b) peptides and (c) NOGs between the two methods. DDA-based: searching the DDA-derived database; DDA-independent: analyzed by MetaLab-DIA workflow. **(d)** the correlation of NOGs between DDA-based and DDA-independent methods. **(e)** the relationship between the number of the peptides and the raw files from different projects.

At the functional level, the DDA-independent approach identified 31,410 NOGs, including 14,090 NOGs not observed in the DDA-based results (Figure 5c**, Supplementary Data 11**). Despite this, the NOGs shared a significant overlap with high correlation (Figure 5d), affirming the comparable credibility of both methods. This dataset serves as a valuable resource for human gut microbiome studies. Compared to all the 15 DDA datasets used to build the MetaPep library, this DIA-PASEF dataset generated the second highest number of peptide identifications (Figure 5e). Project PXD011515 produced approximately 600,000 peptides from over 400 raw files^32^, whereas this DIA-PASEF dataset yielded about 560,000 peptides from just 81 raw files, highlighting the high sensitivity of the DIA mode.

In conclusion, using a DDA-derived database for DIA search limits the power of the DIA strategy, as demonstrated by the less than 300,000 peptides obtained from DDA datasets, restricting DIA peptide identification to 228,746. Our workflow, through deep learning model building and peptide intensity prediction, iteratively generated a comprehensive database containing about 760,000 sequences, yielding approximately 560,000 peptide identifications. This suggests that our workflow outperforms traditional DIA data analysis strategies, simplifying the experimental process and significantly increasing the depth of identification.

## Discussion

The comprehensive analysis of metaproteomics data has been significantly enhanced with the introduction of MetaLab-MAG and MetaLab-DIA. These advanced tools address critical challenges in the field, particularly in the analysis of complex and diverse microbial proteomes with extremely large databases. By integrating innovative data analysis approaches and leveraging the strengths of both DDA and DIA methods, MetaLab-MAG and MetaLab-DIA offer a robust solution for metaproteomic studies.

The MetaLab-DIA software, with its neural network model for predicting peptide candidates, takes full advantage of DIA data by overcoming the limitations of traditional DDA-based library searches. The iterative search strategy and the use of high predicted intensity peptides for DIA searches have demonstrated remarkable improvements in peptide identifications, particularly in complex samples. This approach not only doubled peptide identifications in DIA mode compared to DDA mode but also eliminated the need for additional DDA experiments, thereby streamlining the experimental process.

An essential goal in metaproteomics data analysis is to derive taxonomic information from identified peptides. However, this information requires careful evaluation because the confidence level is typically controlled at the peptide level. For instance, a 1% false discovery rate (FDR) at the peptide level will likely result in a higher FDR at the species level. To address this, MetaLab implemented FDR control at both the genome and taxonomic levels. Specifically, we computed a score for each genome, incorporating the posterior error probability (PEP) and the intensity of all peptides mapped to that genome. Subsequently, a p-value was assigned to each genome, derived from a t-test comparing the score of the target genome with those of decoy genomes. This approach enables us to effectively distinguish false positives in genome identification and provides a robust scoring method to assess the confidence of target identifications.

Our analysis of various datasets, including synthetic microbial communities, mouse gut microbiome samples, and large-scale human gut microbiome datasets from the timsTOF platform, has shown that MetaLab-DIA achieves deeper and more comprehensive proteome coverage. The high sensitivity of DIA mode, coupled with the iterative search strategy, significantly reduces missing values and enhances quantitative analysis consistency. In conclusion, the MetaLab software suite, encompassing MetaLab-MAG and MetaLab-DIA, transcends conventional limitations by supporting diverse mass spectrometry platforms and introducing innovative data analysis approaches. These tools enhance the depth and accuracy of metaproteomic analyses, offering researchers a comprehensive solution to the challenges posed by the complexity and diversity of microbiomes. This advancement adds to the toolbox, not only improve our understanding of microbial functional dynamics but also holding significant potential for applications in human health.

## Methods

### Sample preparation and MS data acquisition

All the raw files used in this paper were downloaded from existing projects. The 12mix and human fecal samples analyzed by both DDA and DIA were from a previous research study by Aakko et al^22^. The timsTOF data from mouse gut microbial samples was from a previous research study by Wang et al^23^. The sample preparation and MS data acquisition about the 21 replicates of DDA and DIA datasets from the human gut microbial samples were described in a previous research study by Duan et al^24^. The high-throughput DIA-PASEF dataset from the human gut microbial samples was from a previous research study by Sun et al^31^.

### Genome and Taxa score

First a score for each peptide was calculated based on its intensity and PEP value across all the raw files.

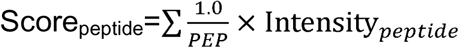

It then assigns these peptides to genomes based on the proteins they match.

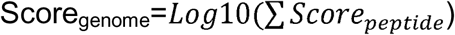

By comparing the intensity of peptides associated with different genomes to decoy genomes, a p-value score for each genome was calculated by t-test.

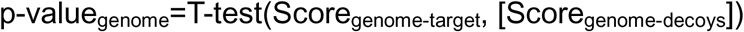

This score represents the confidence level of the genome assignment, with lower scores represent significant different with the decoy genomes, indicating higher confidence. This approach helps to differentiate between true and false positive genome identifications, providing a more reliable assessment of the genome composition of the sample (Supplementary note).

### Predict peptide intensity by deep learning model

This deep learning model utilizes a deep neural network architecture to predict the intensity of peptides based on their underlying features. First a training data table was generated, including peptide sequence, peptide intensity and 15 features: detectability; protein intensity; genome intensity; taxa intensity at seven levels and function intensity at 5 levels. Then the training data table was split into two parts: training sets and testing sets. It also calculates weights for each data point based on the intensity values. Higher intensity peptides get higher weights, making the model pay more attention to them during training. After that the training set was reshaped to a specific format suitable for LSTM layers. Bidirectional LSTMs (Long Short-Term Memory networks) was used to capture sequential information from the peptide features. These layers are capable of learning long-term dependencies within the data. Dense layers with ReLU (Rectified Linear Unit) activation were added for non-linear transformations and Dropout layers were used to prevent overfitting. The model is compiled using the Adam optimizer with a learning rate (also a hyperparameter) and a loss function (mean squared error in this case). It also specifies metrics to monitor during training, including mean squared error, mean absolute error, and weighted mean squared error. A RandomSearch algorithm from Keras Tuner was used to find the optimal hyperparameters for the model. This involves trying out different combinations of hyperparameter values, like number of hidden units in Dense layers and learning rate and selecting the model that performs best on the validation set. A portion of the training data held out for evaluation during hyperparameter tuning. The model was trained with different hyperparameter configurations for a 50 epochs. It also uses early stopping to prevent overfitting. Early stopping monitors the validation loss and stops training if the loss doesn’t improve for a certain number of epochs. After hyperparameter tuning, the best performing model was retrieved and evaluated on the test data. Various performance metrics like mean squared error (MSE), root mean squared error (RMSE), mean absolute error (MAE), and R-squared (R2) were calculated to assess the model’s generalization ability. The best performing model was saved. Finally, it predicted the intensity for peptides from all the related genomes and saves the predicted intensities along with the original peptide sequences. The predicted dataset was served as the source of the peptide candidates. In the iterative search, peptides with higher intensities will be selected to build a peptide library for the peptide identification.

## Data availability

The raw files of 12mix and human fecal samples from Aakko’s paper can be found from ProteomeXchange^33^ with the dataset identifier PXD008738. The raw data of the 21 technical replicates of DDA and DIA analyzed human gut microbiome samples have been deposited to the ProteomeXchange Consortium via the PRIDE partner repository with the dataset identifier PXD056256. The raw files of the DDA-PASEF and DIA-PASEF analyzed mouse gut microbiome samples can be found from ProteomeXchange with the dataset identifier PXD049086 and PXD049089. The human gut microbiome dataset from the timsTOF platform can be found from ProteomeXchange with the dataset identifier PXD051104.

## Software availability

The software tool MetaLab-MAG 1.0 and MetaLab-DIA 1.0 can be downloaded from https://imetalab.ca/.

## Acknowledgement

Funding for this project was provided by the Natural Sciences and Engineering Research Council of Canada (NSERC), and the University of Ottawa. H.D. acknowledges a scholarship from the TECHNOMISE NSERC-CREATE training program.

## Author contributions

K.C. conceptualized and designed the study, developed the software and performed data analysis. Z.N., X.Z., H.D. and J.M. performed data acquisition. K.C. wrote the paper and all authors contributed to the revision. D.F. supervised the overall study.

## Supplementary note

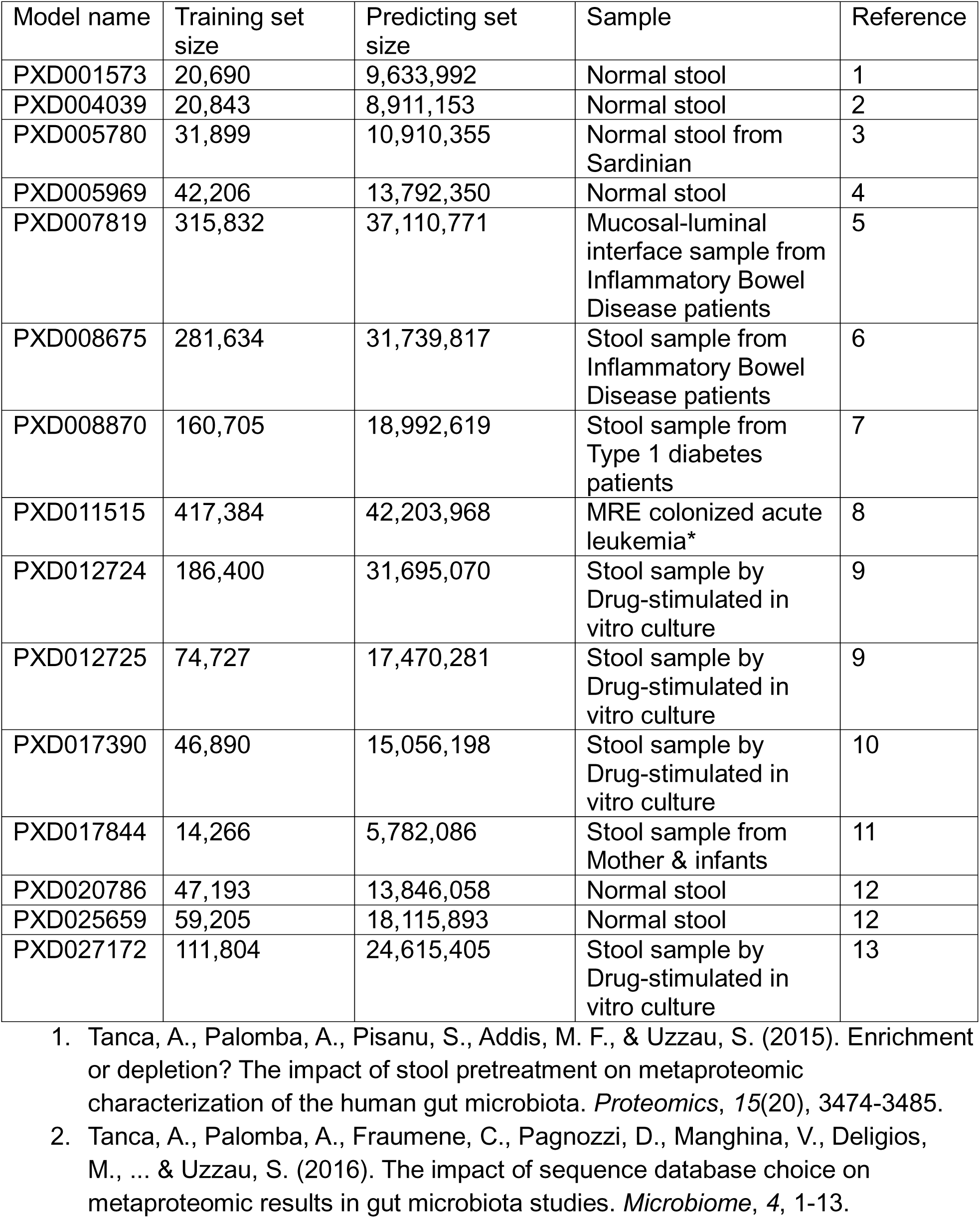
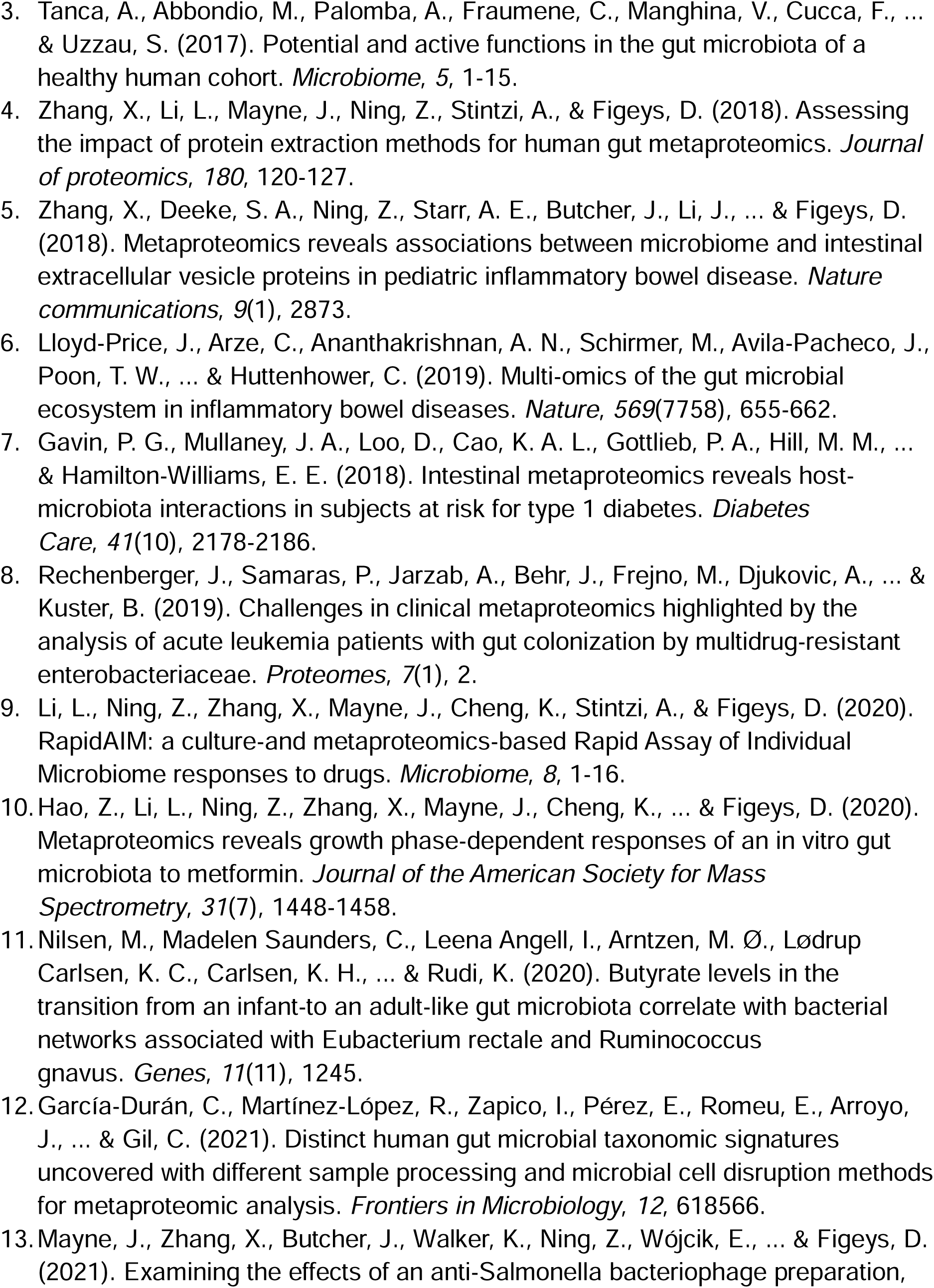

---

**Supplementary figure 1.**
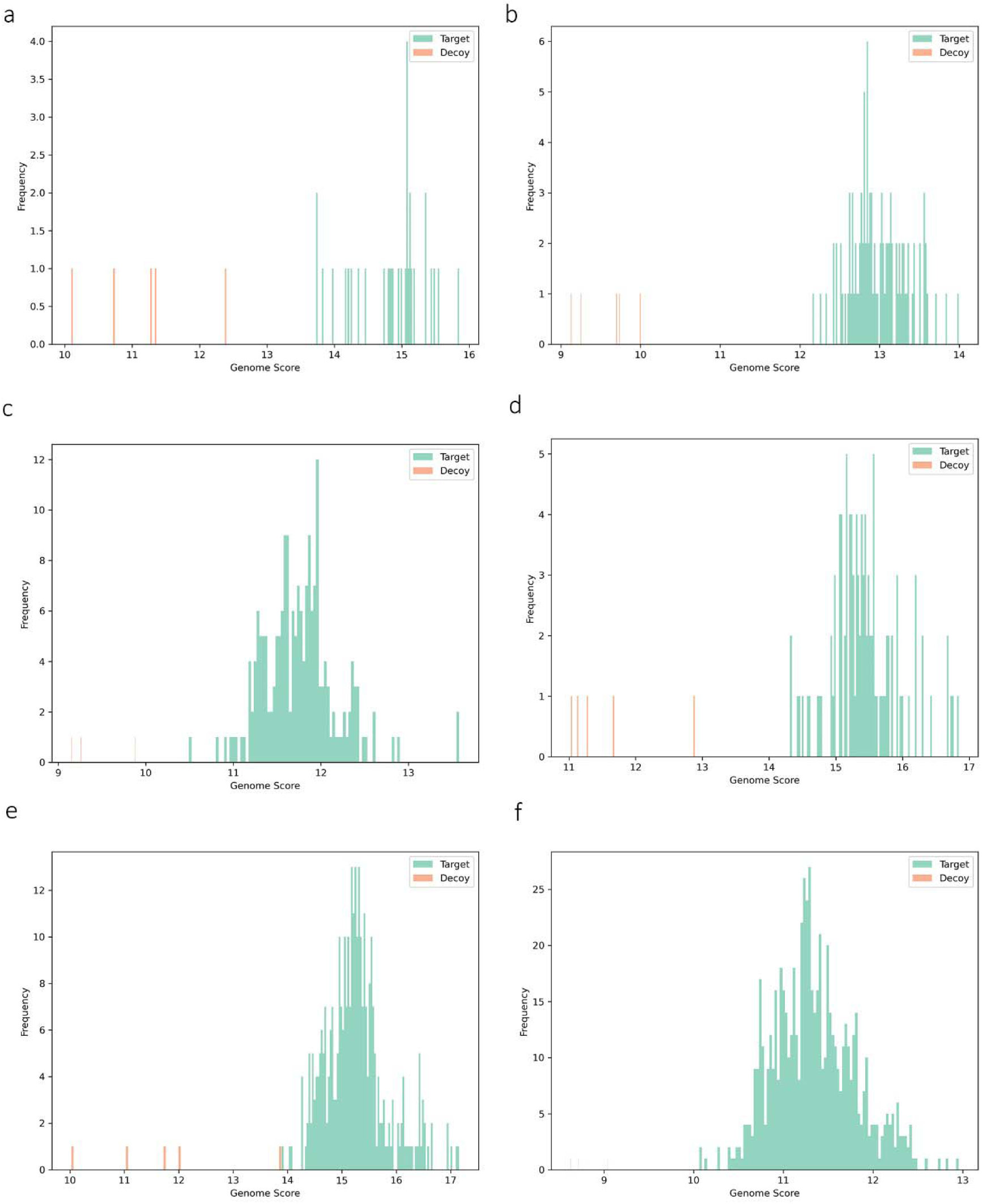
The distribution of the genome scores. (a). the 12mix samples; (b) the fecal samples from the same study of the 12mix samples; (c) the mouse samples analyzed by PASEF-DDA; (d) 21 duplicates human gut microbial samples (DDA-based method); (e) 21 duplicates human gut microbial samples (DDA-independent method); (f) the high-throughput DIA-PASEF samples.

## References

1. Allan Konopka, What is microbial community ecology?, The ISME Journal, Volume 3, Issue 11, November 2009, Pages 1223–1230.

2. Wang, B., Yao, M., Lv, L., Ling, Z., & Li, L. (2017). The human microbiota in health and disease. Engineering, 3(1), 71–82.

3. Ramakrishna, B. S. (2013). Role of the gut microbiota in human nutrition and metabolism. Journal of gastroenterology and hepatology, 28, 9–17.

4. Paul W, Philip L. Bond. (2006) Metaproteomics: studying functional gene expression in microbial ecosystems, TRENDS in Microbiology, 14, 92–97

5. Thilo M, Dirk B, Udo R, Erdmann R and Lennart M. (2013) Searching for a needle in a stack of needles: challenges in metaproteomics data analysis, Molecular BioSystems, 9, 578–585

6. Hu A, Noble WS, Wolf-Yadlin A. (2016) Technical advances in proteomics: new developments in data-independent acquisition. F1000Res. 5: F1000 Faculty Rev-419

7. Zhang, F., Ge, W., Ruan, G., Cai, X., & Guo, T. (2020). Data-independent acquisition mass spectrometry-based proteomics and software tools: a glimpse in 2020. Proteomics, 20(17-18)

8. Pietilä, S., Suomi, T., & Elo, L. L. (2022). Introducing untargeted data-independent acquisition for metaproteomics of complex microbial samples. ISME communications, 2(1), 51.

9. Tsou, C. C., Avtonomov, D., Larsen, B., Tucholska, M., Choi, H., Gingras, A. C., & Nesvizhskii, A. I. (2015). DIA-Umpire: comprehensive computational framework for data-independent acquisition proteomics. Nature methods, 12(3), 258–264.

10. Demichev, V., Messner, C. B., Vernardis, S. I., Lilley, K. S., & Ralser, M. (2020). DIA-NN: neural networks and interference correction enable deep proteome coverage in high throughput. Nature methods, 17(1), 41–44.

11. Cheng, K., Ning, Z., Zhang, X., Li, L., Liao, B., Mayne, J., Stintzi, A. & Figeys, D. (2017). MetaLab: an automated pipeline for metaproteomic data analysis. Microbiome, 5, 1–10.

12. Cheng, K., Ning, Z., Li, L., Zhang, X., Serrana, J. M., Mayne, J., & Figeys, D. (2022). MetaLab-MAG: A metaproteomic data analysis platform for genome-level characterization of microbiomes from the metagenome-assembled genomes database. Journal of Proteome Research, 22(2), 387–398.

13. Li, J., Jia, H., Cai, X., Zhong, H., Feng, Q., Sunagawa, S., et al. (2014). An integrated catalog of reference genes in the human gut microbiome. Nature biotechnology, 32(8), 834–841.

14. Almeida, A., Nayfach, S., Boland, M., Strozzi, F., Beracochea, M., Shi, Z. J., et al. (2021). A unified catalog of 204,938 reference genomes from the human gut microbiome. Nature biotechnology, 39(1), 105–114.

15. Chi, H., Liu, C., Yang, H., Zeng, W. F., Wu, L., Zhou, W. J., et al. (2018). Comprehensive identification of peptides in tandem mass spectra using an efficient open search engine. Nature biotechnology, 36(11), 1059–1061.

16. Yu, F., Teo, G. C., Kong, A. T., Haynes, S. E., Avtonomov, D. M., Geiszler, D. J., & Nesvizhskii, A. I. (2020). Identification of modified peptides using localization-aware open search. Nature communications, 11(1), 4065.

17. Yu, F., Haynes, S. E., Teo, G. C., Avtonomov, D. M., Polasky, D. A., & Nesvizhskii, A. I. (2020). Fast quantitative analysis of timsTOF PASEF data with MSFragger and IonQuant. Molecular & Cellular Proteomics, 19(9), 1575–1585.

18. Strauss, M. T., Bludau, I., Zeng, W. F., Voytik, E., Ammar, C., Schessner, J. P., et al. (2024). AlphaPept: a modern and open framework for MS-based proteomics. Nature Communications, 15(1), 2168.

19. Yang, J., Cheng, Z., Gong, F., & Fu, Y. (2023). DeepDetect: deep learning of peptide detectability enhanced by peptide digestibility and its application to DIA library reduction. Analytical Chemistry, 95(15), 6235–6243.

20. Gurbich, T. A., Almeida, A., Beracochea, M., Burdett, T., Burgin, J., Cochrane, G., et al. (2023). MGnify genomes: a resource for biome-specific microbial genome catalogues. Journal of Molecular Biology, 435(14), 168016.

21. Richardson, L., Allen, B., Baldi, G., Beracochea, M., Bileschi, M. L., Burdett, T., et al. (2023). MGnify: the microbiome sequence data analysis resource in 2023. Nucleic Acids Research, 51(D1), D753–D759.

22. Aakko, J., Pietila, S., Suomi, T., Mahmoudian, M., Toivonen, R., Kouvonen, P., et al. (2019). Data-independent acquisition mass spectrometry in metaproteomics of gut microbiota—implementation and computational analysis. Journal of proteome research, 19(1), 432–436.

23. Wang, A., Fekete, E. E., Creskey, M., Cheng, K., Ning, Z., Pfeifle, A., et al. (2024). Assessing fecal metaproteomics workflow and small protein recovery using DDA and DIA PASEF mass spectrometry. Microbiome research reports. 2024; 3(3):39.

24. Duan, H., Ning, Z., Sun, Z., Guo, T., Sun, Y., & Figeys, D. (2024). MetaDIA: A Novel Database Reduction Strategy for DIA Human Gut Metaproteomics. bioRxiv, 2024–03.

25. Sun, Z., Ning, Z., Cheng, K., Duan, H., Wu, Q., Mayne, J., & Figeys, D. (2023). MetaPep: A core peptide database for faster human gut metaproteomics database searches. Computational and Structural Biotechnology Journal, 21, 4228–4237.

26. Zhang, X., Li, L., Mayne, J., Ning, Z., Stintzi, A., & Figeys, D. (2018). Assessing the impact of protein extraction methods for human gut metaproteomics. Journal of proteomics, 180, 120–127.

27. Li, L., Ning, Z., Zhang, X., Mayne, J., Cheng, K., Stintzi, A., & Figeys, D. (2020). RapidAIM: a culture-and metaproteomics-based Rapid Assay of Individual Microbiome responses to drugs. Microbiome, 8, 1–16.

28. Tanca, A., Palomba, A., Pisanu, S., Addis, M. F., & Uzzau, S. (2015). Enrichment or depletion? The impact of stool pretreatment on metaproteomic characterization of the human gut microbiota. Proteomics, 15(20), 3474–3485.

29. Tanca, A., Palomba, A., Fraumene, C., Pagnozzi, D., Manghina, V., Deligios, M., et al. (2016). The impact of sequence database choice on metaproteomic results in gut microbiota studies. Microbiome, 4, 1–13.

30. García-Durán, C., Martínez-López, R., Zapico, I., Pérez, E., Romeu, E., Arroyo, J., et al. (2021). Distinct human gut microbial taxonomic signatures uncovered with different sample processing and microbial cell disruption methods for metaproteomic analysis. Frontiers in Microbiology, 12, 618566.

31. Sun, Y., Xing, Z., Liang, S., Miao, Z., Zhuo, L. B., Jiang, W., et al. (2024). metaExpertPro: a computational workflow for metaproteomics spectral library construction and data-independent acquisition mass spectrometry data analysis. Molecular & Cellular Proteomics, 100840.

32. Rechenberger, J., Samaras, P., Jarzab, A., Behr, J., Frejno, M., Djukovic, A., et al. (2019). Challenges in clinical metaproteomics highlighted by the analysis of acute leukemia patients with gut colonization by multidrug-resistant enterobacteriaceae. Proteomes, 7(1), 2.

33. Perez-Riverol, Y., Bai, J., Bandla, C., García-Seisdedos, D., Hewapathirana, S., Kamatchinathan, S., et al. (2022). The PRIDE database resources in 2022: a hub for mass spectrometry-based proteomics evidences. Nucleic acids research, 50(D1), D543–D552.

